# An analysis of Species Conservation Action Plans in Guinea

**DOI:** 10.1101/2020.01.27.920751

**Authors:** Charlotte Couch, Denise Molmou, Sékou Magassouba, Saïdou Doumbouya, Mamadou Diawara, Muhammad Yaya Diallo, Sékou Moussa Keita, Falaye Koné, Mahamadou Cellou Diallo, Sékou Kourouma, Mamadou Bella Diallo, Mamady Sayba Keita, Aboubacar Oulare, Iain Darbyshire, Eimear Nic Lughadha, Xander Van Der Burgt, Isabel Larridon, Martin Cheek

## Abstract

To achieve conservation success, we need to support the recovery of threatened species. Yet, <5% of plant species listed as threatened on the IUCN Red List have Species Conservation Action Plans (CAPs). If we are to move from a Red List to a Green List for threatened plant species, CAPs need to be devised and implemented. Guinea is one of the most botanically diverse countries in West Africa. Recent research found that nearly 4000 vascular plants occur in Guinea, a 30% increase from previous estimates. 273 of these plant species are now assessed as threatened with global extinction. There is increasing pressure on the environment from the extractive industry and a growing population. In parallel with implementation of an Important Plant Area programme in Guinea, CAPs were developed for 20 threatened plant species. These plans elaborate conservation efforts needed first to safeguard threatened species both *in situ* and *ex situ* and then to support their recovery. We document the approach used to assemble the Species Conservation Action Plans, and we discuss the importance of having up to date field information, IUCN Red List assessments, and use of a collaborative approach. The need for these plans is increasingly important with recent calculations suggesting a third of African plants are threatened with extinction. This paper outlines initial detailed plant conservation planning in Guinea and offers a template for conservation practitioners in other tropical African countries to follow.

## Introduction

The goal of conservationists is to protect globally threatened species and achieve success through species recovery, eventually recording this on the Green List (Akcakaya et al. 2018). Yet, of the 15,774 threatened plant species treated on the global Red List, only 753 (4.78 %) are reported to have Species Conservation Action Plans (CAPs) in place (IUCN 2019). To help address this massive deficit, we offer an approach to developing CAPs for threatened plant species which has succeeded in Guinea (West Africa), as a template for conservation practitioners in other Tropical African countries.

Guinea is one of the most botanically diverse countries in West Africa. It has nearly 4000 vascular plant species (G. Gosline et al. unpubl. data), a significant increase from the c. 3000 listed in the *Flore (Angiospermes) de la République de Guinée* by Lisowski (2009). This increase reflects an extensive searches over the last five years to inventory the flora of Guinea through the digitisation and georeferencing of historical herbarium records (Magassouba et al. 2014, GBIF 2019) complemented by targeted field expeditions to understudied areas of Guinea by the National Herbarium of Guinea (HNG) and the Royal Botanic Gardens, Kew (RBGK) (Cheek et al. 2018a). As a result of efforts to find additional localities for rare species, we determined at least one likely extinction (*Inversodicraea pygmaea* G.Taylor) due to a hydroelectric dam (Cheek 2018c). A total of 74 published endemic species have been recorded for Guinea, all of which are threatened (G. Gosline et al. unpubl. data, Rokni 2017, 2018, Larridon 2018). Recent estimates of endemism for Guinea ranged from 2.6% (Lisowski, based on a species list) to 4.7% (Sosef et al. 2017 inferred from the RAINBIO dataset. The number of endemic species is set to increase with recent discoveries; several descriptions of new species to science are in progress. The comparatively high plant diversity in Guinea is in part due to the highland areas found in the central and eastern parts of the country. The central Fouta Djallon highlands have many endemic plant species found in a variety of habitats such as sandstone cliffs, sandstone and lateritic bowal (treeless hardpan), and submontane forest (Couch et al. 2019a). However, over past centuries this area has undergone a dramatic change with the expansion of cattle ranging and development of agricultural systems. The submontane forest has become extremely degraded and intact submontane forest has practically disappeared over all of the Fouta Djallon. A recent three year project to identify Tropical Important Plant Areas in Guinea (Couch et al. 2019a) showed that of 35 threatened species not seen for 60 years or more, and not rediscovered during the project, 25 are globally endemic to Guinea and the majority occur in the Fouta Djallon. The Fouta Djallon historically shared some species with the mountain chains of Simandou and Nimba in eastern Guinea, such as *Habenaria jaegeri* Summerh. and *Kotschya lutea* (Portères) Hepper. Both species are likely to be locally extinct in the Fouta Djallon, as they have not been seen despite targeted searches there for 57 and 82 years respectively.

The Tropical Important Plant Areas (TIPAs) of Guinea project recently identified nine threatened habitats and 22 TIPAs (Couch et al. 2019a, Couch et al, 2017). TIPAs are assessed using three criteria: the presence of (1) Threatened species, (2) Botanical richness (including socio-economic species), and (3) Threatened habitats (Darbyshire et al. 2017). Each TIPA assessment also documents past, present and future threats as well as the current protection given to the proposed TIPA site. A variety of threats imperil the flora of Guinea, not least from the mining sector. Guinea is one of the leading exporters of bauxite, producing 95% of African bauxite and 15% of the global share, based on tonnes exported (Alcircle, 2018). It also has considerable reserves of iron ore, gold and diamonds, with smaller reserves of other minerals including nickel, copper, cobalt, manganese and uranium (Guinean Ministry of Mines and Geology, 2016). However, for many of the habitats, in particular the lowland evergreen forest and submontane forest (including gallery forest), the main threats are the unsustainable slash and burn agricultural practices and the cutting of wood for charcoal. A study by Sayer et al. (1992) documented that 96% of original forest had already disappeared from Guinea. In the bowal areas, the main threats are from cattle-ranging and linked management practices causing increases in fire frequency, and from housing development.

New large-scale projects, for example in the mining sector and for hydroelectric power, require companies to carry out detailed Social and Environmental Impact Assessments (SEIAs). These studies should highlight threatened plant species in their concessions. Until now, only 7% of all plant species have been assessed globally for the IUCN Red List (Bachman et al. 2019), including c. 5% of the Guinean flora, making it difficult for environmentalists to demonstrate that protection or mitigation is needed. A GBIF-BID funded project (reference number AF2015-0042-NAC) entitled ‘Towards a Red Data Book for Guinea’ in collaboration with the Darwin Initiative funded TIPAs project (Darwin Project 23-002), has assessed c. 200 plant species from Guinea (using the IUCN 2012 guidelines). This is a considerable achievement. However, the review and publishing process of IUCN Red List assessments are time-consuming. These delays were accentuated because until recently there was no IUCN specialist group available to review most West African plant assessments. In June 2019, the West Africa Plant Red List Authority (WAPRLA) was accepted by the IUCN Species Survival Commission. It is expected that the new RLA will reduce the delays in publication of IUCN Red List assessments of West African plants and will also unite plant conservation efforts and promote red listing across the West African region. A preliminary list of threatened plant species for Guinea was published in PeerJ Preprints in 2017, and updated over the course of the projects described above (Couch et al. 2019b) to keep conservation practitioners up to date ahead of the publication of a full plant Red Data Book for Guinea 2020. This will also feed into updating the *Monographie Nationale* (Guinea’s National Biodiversity Management Action Plan) which has not been revised since 1997. As part of the red listing process, ongoing and required conservation actions are recorded. However, these are high level actions with little or no detail generally given.

As part of the GBIF-BID ‘Towards a Red Data Book for Guinea’ project, and as a first step towards detailed plant conservation planning in Guinea, individual Species Conservation Action Plans were developed for 20 plant species assessed as Critically Endangered (CR), Endangered (EN) or Vulnerable (VU). These plans document the conservation efforts needed to safeguard each of these threatened species both *in situ* and *ex situ*.

In this paper, we outline the approach used to assemble the Species Conservation Action Plans, and we discuss the importance of having up to date species field information and IUCN Red List assessments. We also discuss the advantages of a collaborative approach and outline the next stages for implementing these species action plans on the ground in Guinea.

## Materials and Methods

Several points need to be considered before a Species Conservation Action Plan can be written. Firstly, who should be involved in this process? To address this and the assessment of the Tropical Important Plant Areas, a joint working group on TIPAs and Conservation Action Plans (CAPs) was formed in May 2018. The working group consists of representatives from the National Herbarium of Guinea (HNG), the Royal Botanic Gardens, Kew, UK (RBGK), the Guinean Ministry of Environment, Water and Forests (MEEF), the National Parks and Reserves office (MEEF-OGuiPAR), the Centre for Biological Observations and Monitoring (MEEF-COSIE), environmental NGOs Guinée Ecologie (GE) and Protection et Gestion de l’Environnement (PEG), the Centre for Environmental Research Studies at the Université Gamal Abdel Nasser de Conakry (UGANC-CERE), and the Seredou Herbarium (Institut de Recherche Agronomique de Guinée (IRAG) acronym SERG). This was the first time that these organisations had united to support the prioritisation of threatened plant conservation in Guinea. The working language of the group is French. The working group meets every 2 months and conducts business over email inbetween meetings, discusses and agrees what should be included in the CAPs and which designated members are to be charged with collating the information and writing the plans.

The protocol for preparing the conservation action plans was developed and approved by the working group (available on the National Herbarium of Guinea website) and was based in part on the *Conservation Action Planning Handbook* by The Nature Conservancy (TNC 2007). The format and style of the Conservation Action Plans drew upon previously drafted species recovery plans for non-Guinean taxa (e.g. JNCC UK priority species pages 2010a, 2010b, Panjabi et al. 2011) together with constent from conservation actions identified in the IUCN Red List process.

A shortlist was drawn up by the HNG and RBGK members of the group from the preliminary list of threatened species of Guinea (Couch et al. 2019b). Twenty species were chosen that meet the following selection criteria: 1) listed as CR, EN or VU in the preliminary checklist, 2) have a published or reviewed IUCN Red List assessment, 3) cover a range of life forms, and 4) are found over a range of threatened habitats. The decision to prepare CAPs only for species with formal IUCN assessments was made in part because there is more information available for these species, but also because their assessed conservation status was unlikely to change in the near future. With species that have yet to be formally assessed or reviewed there is the risk that the IUCN Red List status may change. The species chosen included a mixture of life forms i.e. trees, shrubs, lianas and herbs (Table 1).

**Table 1.** List of species chosen for Conservation Action Plans. IUCN status: CR = Critically Endangered, EN = Endangered, VU = Vulnerable.

| Family | Species | IUCN status | Growth form |
| --- | --- | --- | --- |
| Acanthaceae | *Anisotes guineensis* Lindau | EN | Shrub |
| Apocynaceae | *Marsdenia exellii* C.Norman | EN | Liana |
| Apocynaceae | *Xysmalobium samoritourei* Goyder | EN | Herb |
| Araceae | *Stylochaeton pilosus* Bogner | EN | Herb |
| Asteraceae | *Vernonia djalonensis* A.Chev. | CR | Herb |
| Bromeliaceae | *Pitcairnia feliciana* (A.Chev.) Harms & Mildbr. | CR | Herb |
| Cyperaceae | *Scleria guineensis* J.Raynal | CR | Herb |
| Ebenaceae | *Diospyros feliciana* Letouzey & F.White | EN | Tree |
| Euphorbiaceae | *Acalypha guineensis* J.K.Morton & G.A.Levin | VU | Herb |
| Lamiaceae | *Plectranthus linearifolius* (J.K.Morton) B.J.Pollard & A.J.Paton | EN | Herb |
| Leguminosae-Detarioideae | *Talbotiella cheekii* Burgt | EN | Tree |
| Leguminosae-Papilionoideae | *Pterocarpus erinaceus* (DC.) Polhill & Wiens | EN | Tree |
| Leguminosae-Papilionoideae | *Eriosema triformum* Burgt | CR | Herb |
| Melastomataceae | *Cailliella praerupticola* Jacq.-Fél. | EN | Shrub |
| Orchidaceae | *Habenaria jaegeri* Summerh. | EN | Herb |
| Podostemaceae | *Inversodicraea pepehabai* Cheek | EN | Herb |
| Rubiaceae | *Keetia susu* Cheek | EN | Shrub/Tree |
| Rubiaceae | *Tarenna hutchinsonii* Bremek. | CR | Shrub/Tree |
| Rutaceae | *Vepris felicis* Breteler | CR | Shrub |
| Sapotaceae | *Tieghemella heckelii* (A.Chev.) Pierre ex Dubard | EN | Tree |

With the protocol drafted and the short list of species agreed, two members of the group took the lead on collating species information and drafting the plans. All members of the group contributed to review and refinement of the plans. This work took place over a period of 9 months and involved an estimated 152 person working days.

The first part of each CAP sets out the context for each species. It is imperative that each species is properly researched and clearly circumscribed based on sound taxonomy. This is especially necessary since any existing documentation, particularly in Guinea, is often out of date. Until recently, there was little published on the Guinean flora. The *Flore (Angiospermes) de la Guinée* by Lisowski, was published posthumously in 2009, with taxonomy not updated since Lisowski submitted it for publication in 2000. As a result, many names in Lisowski’s *Flore de la Guinée* are out of date (Cheek et al. 2015), and since its publication about 20 newly discovered species (e.g. Cheek & Haba 2016, Cheek et al. 2018b) and many new range extensions have been recorded in Guinea. Where a recently discovered species was chosen for a CAP, the protologue (original scientific publication) has been used as the source for the taxonomic information. Names have been checked against the African Plant Database (2019), the International Plant Names Index (2019) and Plants of the World Online (2019). Descriptions of the plant species were taken from either Lisowski (2009), the *Flora of West Tropical Africa* (Keay & Hepper 1954-72) or the protologue. Details about the ecology, phenology and habitat where known, have also been documented.

The working group decided that each CAP should include as much information as is available for the species including i) past and present collection data, this information has largely been collated from herbarium specimen label data and species accounts in the works cited above. If the species is known to be used by people, these uses are also documented; ii) geographical distribution, particularly within Guinea to focus conservation efforts and, iii) where known, the number of indviduals in the population. Specimen-based distribution maps for each species have been produced based on records collected for the Red Listing programme, examples can be seen in Fig. 1. Distribution maps were made using ArcGIS Pro software with simple XY coordinates uploaded and mapped onto a world basemap.

**Fig. 1.**
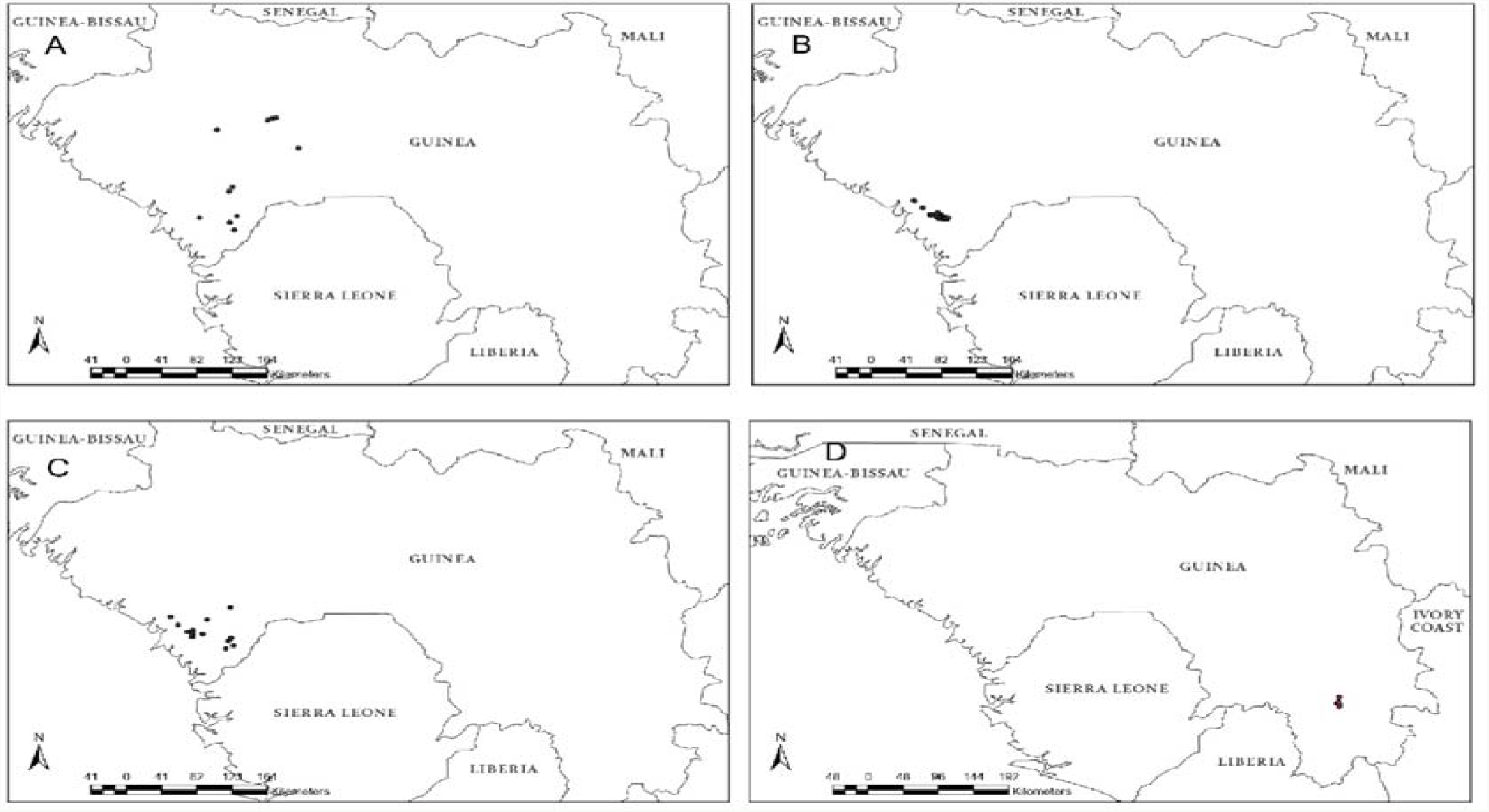
Species distribution maps for four endemic Guinean species. **A** *Anisotes guineensis*, **B** *Talbotiella cheekii*, **C** *Diospyros feliciana*, and **D** *Eriosema triformum*.

Information on threats both past and present, direct and indirect, is listed. This information was gathered partly through literature but also during recent fieldwork. As part of the Darwin Initiative funded project on Tropical Important Plant Areas in Guinea 2016-2019, over 20 field expeditions were carried out targeting rare species and priority threatened habitats. These field expeditions were invaluable to gather current information on rare species, their distribution and uses, and on the current threats.

The second part of the CAP document sets out a summary plan for the management and conservation of the species based on current knowledge. The first section, of the second part, details all known research or suggests what research is required. *In situ* and *ex situ* conservation actions are then proposed for the protection of the species.

*In situ* conservation actions detail any protected areas in which the species is currently found, and whether the species is found within any of the newly designated Tropical Important Plant Areas (Couch et al. 2019a). The total size of the population and details of the sites where the species is found are recorded so that these data can be presented to the local authorities and ultimately support the legal protection of the sites and species. The CAP also emphasises that this documentation process must always be undertaken with support from local communities especially when a species (sub)population occurs within a community forest or sacred forest or in an area outlined for housing development, e.g. as is the case for *Vernonia djalonensis* A.Chev. Without community support, conservation efforts will have little long-term effect on the survival of species on the ground.

*Ex situ* conservation actions focus on the propagation of the species outside its range, seed collection and banking where applicable, and the potential for translocation to another protected area or botanical garden. The results of any experiments previously completed are also documented. Each CAP also recommends sensitization of the local population to the importance of plant species conservation and to the protection of the national plant heritage of Guinea.

## Results

Of the twenty CAPs produced (see Online Resources 1-20), 11 are for species endemic to Guinea; this represents 15% of the total 74 published endemic plant species of Guinea. The threats to the CAP species vary. All species have one or more associated threats. Fig. 2 shows the percentage of species per threat type. The threats affecting most CAP species are uncontrolled fires (75%), mining or quarrying (60%) and infrastructure / urbanisation (50%). Two of the species are directly threatened by pollution and all of the woody species (9) are threatened by deforestation or clearance of habitat through slash and burn agriculture, which is also a threat to 40% of the CAP species overall.

**Fig. 2.**
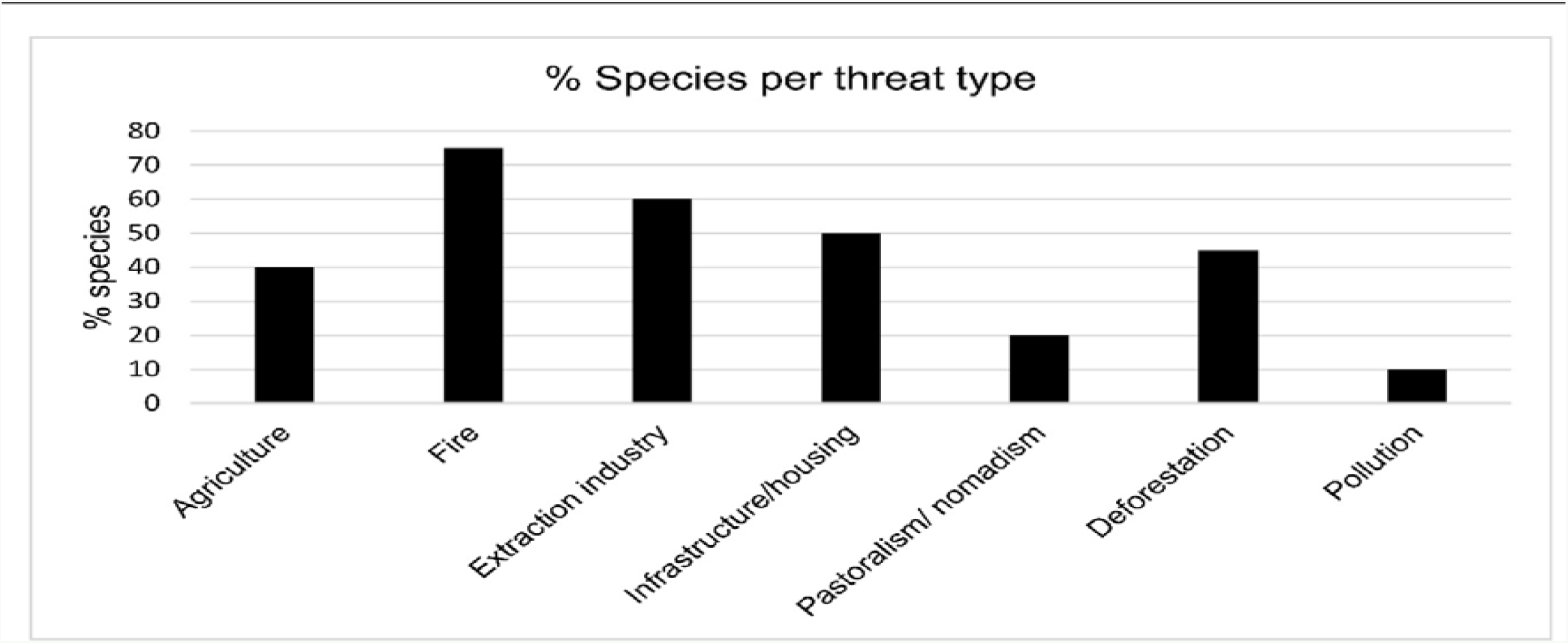
Graph showing the percentage of the 20 Conservation Action Plan species per threat type.

Nine of the twenty CAP species are found in a current protected area and all of the species are found within one or more of the newly designated TIPAs (Couch et al. 2019a). However, these protected areas either lack management plans or have management plans which are outdated. Within these management plans, specific species conservation actions, especially for plants, are usually absent.

Eight of the CAP species have seed collections made and banked at the Simfer base in the Simandou mountains or Herbier National de Guinee, and the Millenium Seed Bank at RBGK, UK, though none have reached the recommended seed banking target threshold of 10,000 seeds (Way & Gold 2014). Some seed collections are small because there are few known individuals or individuals do not produce many seeds each season. Some species have large seeds, expected to be recalcitrant, i.e. they are unsuitable for conventional seed banking, the seeds dying when dried. *Talbotiella cheekii* Burgt is one such species (Burgt et al, 2018). Some Rubiaceae species are also known to be recalcitrant so *Tarenna hutchinsonii* Bremek. and *Keetia susu* Cheek may also prove to be unsuitable for conventional seed banking, but as yet they remain untested.

For the majority of the CAP species no propagation information is available and so experimentation will be required to fill this knowledge gap. However, for a quarter of the species propagation protocols are available, due to their association with a mining project. These protocols were researched at RBGK using a variety of methods, e.g. micropropagation for *Habenaria jaegeri* (Cheek 2017), and cuttings for *Tarenna hutchinsonii* (Cheek *et al* 2015) and *Marsdenia exellii* C.Norman (Cheek 2013).

Currently, only five of the 20 species have been identified as suitable for potential reintroduction. Two transplant experiments have already been carried out with *Eriosema triformum* Burgt. Transplantation of tubers, from the Simandou mountains to Mt Béro in May 2012, was unsuccessful as the tubers were mostly eaten by squirrels and rock hyrax, and ultimately, all died (Cheek et al, 2017). Translocation of *Eriosema triformum* seed to the Mts Nimba Strict Nature Reserve and also to Mt Tibe was attempted in April 2019. Results of these transplants are to be evaluated in 2020 (X. van der Burgt, pers. comm.). Rhizomes of *Stylochaeton pilosus* Bogner were successfully translocated in 2013 (C. Couch, pers. obs.).

## Discussion

The BID-GBIF funded project “Towards a Red Data book for Guinea” and the Darwin Initiative funded project identifying “Tropical Important Plant Areas of Guinea” have the attracted attention of both national and international audiences to the threatened and unique plant species of Guinea. The Species Conservation Action Plans resulting from these projects are the first to be written for threatened plant species in Guinea and are a result of the collaboration between the HNG, RBGK, Guinean government departments and NGOs focussed on plant conservation. The partnerships and expertise on plant conservation built during these projects did not previously exist. Conservation of endemic and near-endemic plant species had not been on the national agenda. In contrast, the conservation of large mammals such as the chimpanzee have had high levels of attention (Sugiyama & Soumah 1988, Brugière et al. 2005, Fleury-Brugière & Brugière 2010, Humle et al. 2011). Following the conclusion of the Darwin and GBIF-BID projects in March 2019, the working group has continued to collaborate to address, review and update 1) Guinea’s CITES list and 2) the second edition of the Guinean National Biodiversity Action Plan. The collaboration has gained support from all sectors concerned with plant conservation and together with the recently identified Tropical Important Plant Areas (Couch et al. 2019a), it is pushing plant conservation in Guinea further up the national agenda. To the best of our knowledge this is the first time that a programme of plant conservation action plans for globally threatened plant species has been devised and acted upon in a West African country. The 20 Species Conservation Action Plans have highlighted the importance of fieldwork to provide up to date information on the target species. IUCN Red List assessments can be written based on literature and herbarium records, but knowing the current status of a population and the real threats that they face is invaluable when writing a CAP. The CAPs draw on fieldwork undertaken over the past 10 years, largely by HNG and RBGK.

Implementation of these plans will need further action. One of the plans, for *Vernonia djalonensis* (Online Resource 19), was the basis of a successful funding bid to the Mohammed bin Zayed Species Conservation Fund and is being used to guide on-the-ground conservation actions for this species, which was recently elected as the national flower of Guinea (Couch 2018). We have engaged with the local authorities and local plant nurseries to protect and propagate this species. In early 2020, we are planning to engage with students and youth groups.

There is a growing need for documentation on the plants of Guinea, not only because there has been so little published, but also because much of the documentation is out of date. The need for national scale CAPs for individual priority species is ever more important as original habitat is lost due to human activities. One in five of the world’s plant species were reported as threatened (RBG Kew 2016), and Stévart et al. (2019) infer that one in three African plant species are at risk of extinction. With 96% loss of original forest reported in 1992 (Sayer et al. 1992), many of Guinea’s most threatened plant species need tailored CAPs if they are not to follow *Inversodicraea pygmaea* into global extinction.

Guinea has over 270 threatened species of which 74 are endemic. Given the person hours required to write 20 CAPs for threatened species, some endemic species might be more effectively treated by local area action plans, for example based on the 22 identified TIPAs, provided they adequately cover the species encompassed (Monteiro et al. 2018). Broader action plans for example a comprehensive action plan that treats all Guinean threatened tree species might be more effective than individual CAPs. This would reduce the timeframe needed to develop conservation action plans enabling efficient implementation. Increased development in Guinea is resulting in an increase in environmentally damaging projects e.g. mining, hydroelectric dams and quarrying. However, development does not have to mean wholesale destruction of the environment and global extinction of species. Good management based on solid data and analysis can lead to much better industry practices. Guinea’s mining projects are implicated in the conservation of the plant species found in their concessions, but they do not always make this data available. There is a vital need to have up to date information freely available for those assessing the environmental impact of such projects and the possible mitigation that can be achieved. Most of the existing conservation efforts in Guinea are focussed on mammals, birds, ecosystem services, commercial trade or large-scale landscape protection e.g. trans-boundary areas such as the Nimba Mountains (STEWARD 2008, Nganje et al. 2014, Brugière & Kormos 2008, Brugière 2012, Brugière et al. 2005, Correia et al. 2010, Samoura et al. 2007). TIPAs aside, most of the currently protected areas in Guinea do not overlap with concentrations of threatened plant species (Couch et al. 2019a). The majority of the Classified Forests (CF) were designated for forestry services, are not protected within the National Parks and Reserves network, and are considered as unprotected by the Guinea Government. In those few cases where a CF is considered protected, it will have a second designation e.g. ‘Reserve Intégrale’ or National Park. Where management plans are in place for protected areas, these do not include protection of individual threatened plant species or their specific habitats for example the 1995-2014 Management plan for the forest of Ziama (PROGERFOR, 1994).

The 20 species CAPs will be used to target plant conservation and funding, not just in protected areas. They also have the potential to form the basis of conservation planning and mitigation strategies for the extractive industries in those cases where project footprints intersect with those of the threatened plant species. With 20 species CAPs written of 273 threatened species in Guinea, this is merely the beginning. On the ground implementation of the 20 CAPs will assist in updating and modifying the CAP protocol to make it a useful and relevant tool in future conservation planning in Guinea.

## Author Contributions

Writing and compilation of supplementary materials: CC and DM; Contribution and revision of CAPs: all members of the working group; Revision of content and supplementary materials: XvdB, IL, MC, EL. Writing of BID funding proposal: ID, MC.

## Acknowledgements

This work was funded through a grant from Global Biodiversity Information Facility (GBIF) Biodiversity Information for Development (BID) programme. Isabel Larridon is supported by the B.A. Krukoff Fund for the Study of African Botany. The authors would like to thank all those involved with field work in Guinea over the past 10 years who have gathered data, notably Pépé Haba, Gbamon Konomou, Pierre Haba, Fatoumata Fofana Madé, Almamy Diallo, Natalie Konig, Oliver Hooper and Abdoulaye Baldé. Nagnouma Conde, Tokpa Seny Dore, Albert Guilavogui, Boubacar Sow, and Saba Rokni. We also thank Royal Botanic Gardens Kew volunteers Margaret Joachim and Rosemary Lomer who georeferenced herbarium specimens for the GBIF-BID and Darwin Initiative Tropical Important Plant Areas Guinea projects. Our thanks also to Catia Canteiro and Emma Williams for their work on writing IUCN Red List assessments for Guinean plant species. This work has been enabled by the Memorandum of Understanding between RBG Kew and National Herbier de Guinee since 2008.

## Ethical standards

This research was carried out in accordance with the *Oryx* code of conduct.

## Supplementary files

**Conservation Action Plans for 20 threatened Guinean plant species can be found following the links below:**

CAP 1. *Acalypha guineensis* J.K.Morton & G.A.Levin DOI: 10.13140/RG.2.2.34363.36648
CAP 2. *Anisotes guineensis* Lindau DOI: 10.13140/RG.2.2.25974.75845
CAP 3. *Cailliella praerupticola* Jacq.-Fél. DOI: 10.13140/RG.2.2.34363.36648
CAP 4. *Diospyros feliciana* Letouzey & F.White DOI: 10.13140/RG.2.2.15824.87047
CAP 5. *Eriosema triformum* Burgt DOI: 10.13140/RG.2.2.35957.5296
CAP 6. *Habenaria jaegeri* Summerh. DOI: 10.13140/RG.2.2.11630.56644
CAP 7. *Inversodicraea pepehabai* Cheek DOI: 10.13140/RG.2.2.25052.33925
CAP 8. *Keetia susu* Cheek DOI: 10.13140/RG.2.2.18341.45280
CAP 9. *Marsdenia exellii* C.Norman DOI: 10.13140/RG.2.2.28407.78244
CAP 10. *Pitcairnia feliciana* (A.Chev.) Harms & Mildbr. DOI: 10.13140/RG.2.2.21696.89609
CAP 11. *Plectranthus linearifolius* (J.K.Morton) B.J.Pollard & A.J.Paton DOI: 10.13140/RG.2.2.35118.66880
CAP 12. *Pterocarpus erinaceus* (DC.) Polhill & Wiens DOI: 10.13140/RG.2.2.30085.50401
CAP 13. *Scleria guineensis* J.Raynal DOI: 10.13140/RG.2.2.23374.61767
CAP 14. *Stylochaeton pilosus* Bogner DOI: 10.13140/RG.2.2.36796.39049
CAP 15. *Talbotiella cheekii* Burgt DOI: 10.13140/RG.2.2.30164.14728
CAP 16. *Tarenna hutchinsonii* Bremek. DOI: 10.13140/RG.2.2.20097.81766
CAP 17. *Tieghemella heckelii* (A.Chev.) Pierre ex Dubard DOI: 10.13140/RG.2.2.33519.59047
CAP 18. *Vepris felicis* Breteler DOI: 10.13140/RG.2.2.18420.09606
CAP 19. *Vernonia djalonensis* A.Chev. DOI: 10.13140/RG.2.2.15064.65289
CAP 20. *Xysmalobium samoritourei* Goyder DOI: 10.13140/RG.2.2.28486.42561

